# Homeostatic Synaptic Scaling Establishes the Specificity of an Associative Memory

**DOI:** 10.1101/2020.12.04.412163

**Authors:** Chi-Hong Wu, Raul Ramos, Donald B Katz, Gina G Turrigiano

## Abstract

Accurate memory formation has been hypothesized to depend on both rapid Hebbian plasticity for initial encoding, and slower homeostatic mechanisms that prevent runaway excitation and subsequent loss of memory specificity. Here, we tested the role of synaptic scaling in shaping the specificity of conditioned taste aversion (CTA) memory, a Hebbian plasticity-dependent form of associative learning. We found that CTA memory initially generalized to non-conditioned tastants (generalized aversion), becoming specific to the conditioned tastant only over the course of many hours. Blocking synaptic scaling in the gustatory cortex (GC) prolonged the duration of the initial generalized aversion and enhanced the persistence of synaptic strength increases observed after CTA. Taken together, these findings demonstrate that synaptic scaling is important for sculpting the specificity of an associative memory and suggest that the relative strengths of Hebbian and homeostatic plasticity can modulate the balance between stable memory formation and generalization.

## Introduction

Ethologically relevant learning is a delicate balance between the specific details of a memory and what can be generalized to similar circumstances. Because no scenario will reoccur exactly, a balance between memory specificity and generalization can endow an animal with cognitive flexibility^1^. However, the neural mechanisms that shape the specificity of memory remain poorly understood. The most extensively studied cellular process for the encoding of associative memories is the Hebbian modification of synapses, primarily, long-term potentiation (LTP)^2,3,4^. While LTP is generally considered critical for learning and memory^5,6,7^, it may not be sufficient to faithfully encode memories due to the positive feedback nature of Hebbian learning rules^8,9^. Left unchecked, this positive feedback could give rise to the unconstrained enhancement of synaptic strengths, which in turn might result in the degradation of memory specificity^10^. These theoretical considerations suggest that for a memory to be properly encoded, additional homeostatic plasticity mechanisms are required to counterbalance this “runaway” plasticity.

Homeostatic synaptic scaling is a cell-autonomous, negative feedback mechanism that bidirectionally scales excitatory post-synaptic strengths up or down to maintain neuronal activity within a set-point range^11,12,13,14^. Synaptic scaling is hypothesized to stabilize neural network activity in the face of learning-driven changes in synaptic strength^15,16^, and modeling work suggests that synaptic scaling can interact with LTP to enable stable memory formation^9,17^. However, while Hebbian plasticity is rapidly induced^3,4^, synaptic scaling unfolds over many hours^12,16^, suggesting that it cannot stabilize Hebbian plasticity on shorter timescales^18^. This temporal dissociation suggests that unopposed Hebbian plasticity during the early stage of associative memory formation might result in a more generalized memory; by homeostatically scaling down synaptic weights, synaptic scaling might then establish memory specificity over the course of subsequent hours. While compelling on theoretical grounds, this hypothesis has not been tested and the behavioral consequences of disrupted synaptic scaling on associative memory remain unknown.

Here we directly examined how synaptic scaling shapes memory specificity in conditioned taste aversion (CTA) learning, a form of associative learning that relies on Hebbian plasticity within the gustatory cortex (GC)^19,20,21,22,23^. We found that following CTA conditioning, animals transitioned from a generalized to a taste-specific aversion over a timescale of ∼24 hours. Blocking synaptic scaling in the GC using viral manipulations prolonged this phase of generalized aversion. Additionally, we found that when animals exhibited a generalized aversion, GC neuronal-ensembles active during conditioning were robustly reactivated by the novel tastant. Abolishing synaptic scaling led to a persistent increase in postsynaptic strengths onto neurons in these GC conditioning-active ensembles, that correlated with the prolonged generalized aversion. Our work demonstrates that synaptic scaling is important for sculpting the specificity of an associative memory and that the homeostatic regulation of synaptic strengths is important for establishing the balance between stable memory formation and generalization.

## Results

### CTA memory specificity emerges over a timescale of hours

Following the induction of CTA learning, animals can exhibit a non-specific generalized aversion to tastants other than the conditioned stimulus^24,25^. The timescale over which this generalized aversion transitions to a specific aversion for the conditioned tastant is unknown, as are the plasticity mechanisms that could support this transition. To investigate this, we designed a two-bottle CTA paradigm (Fig. 1a). The two-bottle choice test is a standard CTA protocol with the advantage that it is sensitive to a range of aversion strengths^20^. Young Long-Evans rats (postnatal days p28-p32) of both sexes underwent CTA conditioning using saccharin as the CS. Male and female rats showed comparable CTA and generalized aversions so data from both sexes were combined (Supp. Fig. 1). After tastant presentation, rats were injected with LiCl (0.15 M, Moderate CTA group). A memory test with saccharin (CTA Test) 4 hours post-conditioning revealed an aversion in the Moderate CTA group compared to the CS Only control group (Fig. 1b). When we tested for a generalized aversion (Gen. Test), the Moderate CTA group demonstrated a significant decrease in its preference for NaCl (Fig. 1c), a novel tastant that is easily discriminated from saccharin^26^. This decrease was not a result of past tastant experience or LiCl-induced malaise, as no aversion was evident in the CS Only and US Only controls, respectively (Fig. 1c). These results demonstrate that during the early stage of CTA-memory acquisition, animals exhibit an aversion that is not specific to the conditioned stimulus. We next sought to determine how long this generalized aversion persists by performing the Gen. Test at 24 hours post-conditioning in a separate group of rats. At this timepoint, Moderate CTA rats demonstrated no significant difference in taste preference score compared to CS Only controls (Fig. 1d). These results indicate that, over a time course of many hours, CTA memory transitions from a non-specific generalized aversion to a memory that is specific to the conditioned stimulus.

**Fig. 1.**
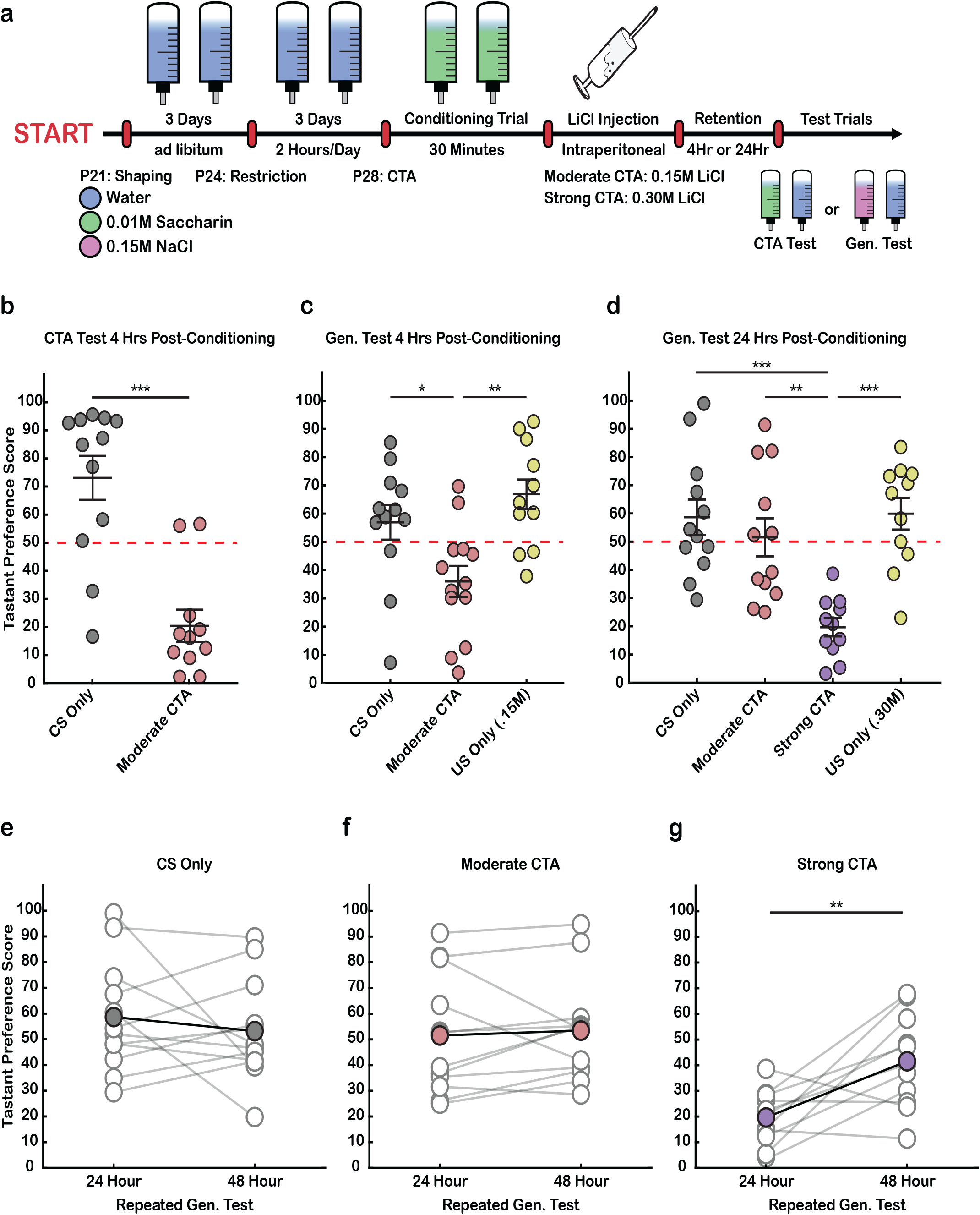
Conditioned Taste Aversion (CTA) memory specificity emerges over a timescale of hours. **a)** Detailed two-bottle CTA learning paradigm; tastant preference scores [(total tastant/total consumed) · 100] were calculated for CTA acquisition (CTA test) or for generalized aversion (Gen. Test) at either 4- or 24-hours post-conditioning. **b)** Moderate CTA; preference for saccharin tested 4Hrs post-conditioning to saccharin (CS Only, n = 12; MCTA, n = 11; Wilcoxon Rank Sum test, p = 0.0005; error bars represent SEM). **c)** Gen. test; preference for NaCl tested 4 hours after moderate CTA conditioning (CS Only, n = 12; MCTA, n = 13; US Only (.15 M LiCl), n = 11; One-way ANOVA, p = 0.0014; Tukey-Kramer post-hoc test, CS Only VS MCTA p = 0.0319, CS Only VS US Only p = 0.4401, US Only VS MCTA p = 0.0012). **d)** Gen. test; preference for NaCl tested 24 hours after moderate or strong CTA conditioning (CS Only, n = 12; MCTA, n = 12; SCTA, n = 11; US Only (.30 M LiCl), n = 11; One-way ANOVA, p = 0.00002; Tukey-Kramer post-hoc test, CS Only VS MCTA p = 0.8029, CS Only VS SCTA p = 0.0001, CS Only VS US Only p = 0.9986, MCTA VS SCTA p = 0.0016, MCTA VS US Only p = 0.7272, SCTA VS US Only p = 0.0001). **e-g)** Gen. test; preference for salt tested 24- & 48-hours post-conditioning (CS Only paired t-test, p = 0.4572; MCTA paired t-test, p = 0.6441; SCTA paired t-test, p = 0.0073). For behavior data here and below, each point represents an individual animal, and the mean and SEM are indicated by line and error bars. Dashed red line indicates no preference.

### The onset of memory specificity depends on the intensity of conditioning

We wanted to understand how the intensity of CTA conditioning influences the duration of the generalized aversion. Previous work has demonstrated that different concentrations of LiCl produce relative differences in CTA strength^27^. Therefore, we wondered whether a stronger conditioning event would trigger a greater cellular response and prolong the duration of the generalized aversion. As opposed to the Moderate CTA group, which exhibits no generalized aversion after 24 hours, the Strong CTA group (.30 M LiCl) exhibited a persistent generalized aversion 24 hours post-conditioning (Fig. 1d). This aversion was not a result of more substantial malaise induced by LiCl treatment, as the US Only (.30 M) controls showed no generalized aversion (Fig. 1d). Thus, stronger CTA conditioning prolonged the duration of the generalized aversion. These experiments reveal an interaction between the strength of conditioning and the onset of CTA memory specificity. In order to better understand this process, we performed repeated testing to characterize the attenuation of the generalized aversion induced by stronger CTA conditioning. We observed that while tastant preference in control groups did not change, the generalized aversion induced in the Strong CTA group reversed upon repeated exposure to NaCl (Fig. 1e-g). There were no differences between any experimental conditions in tastant consumption during conditioning (Supp. Fig. 2a-c). Animals in both moderate and strong conditioning groups demonstrated significant aversions to saccharin after generalized aversion testing, confirming the formation of CTA memory (Supp. Fig. 2d,e). Notably, but not surprisingly, conditioning resulted in a floor effect across both moderate and strong groups due to the sensitivity of the two-bottle choice test (see Bures et al., 1998 for an exploration of CTA methodology). Together, these results demonstrate that the time course over which the specificity of CTA memory emerges is sensitive to the intensity of CTA conditioning.

**Fig. 2.**
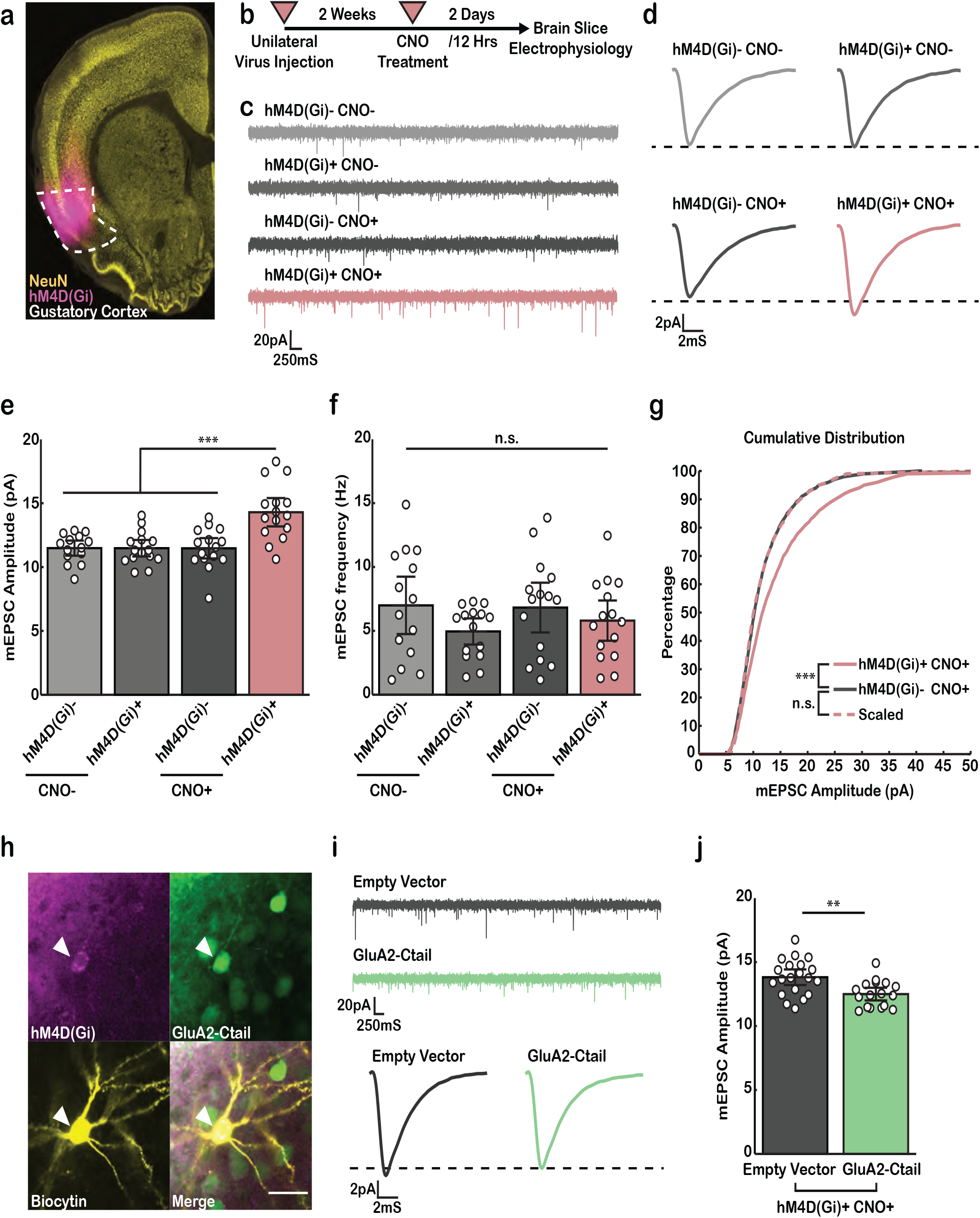
Neurons in the gustatory cortex express homeostatic synaptic scaling. **a)** Coronal brain slice containing the gustatory cortex (outlined in white), showing neurons expressing the inhibitory DREADDS hM4D(Gi) (magenta signal). **b)** Experimental protocol. **c)** Representative mEPSC recordings. **d)** Average mEPSC waveform for each condition, black dashed line is aligned to hM4D(Gi)^□^ CNO^□^ waveform peak. **e)** Cell-average mEPSC amplitudes (hM4D(Gi)^□^ CNO^□^, n = 14; hM4D(Gi)^+^ CNO^□^, n = 15; hM4D(Gi)^□^ CNO^+^, n = 15; hM4D(Gi)^+^ CNO^+^, n = 15; One-way ANOVA, p = 5.8849e-06; Tukey-Kramer post-hoc test, hM4D(Gi)^+^ CNO^+^ VS hM4D(Gi) ^□^ CNO^+^ p = 0.0001, hM4D(Gi)^+^ CNO^+^ VS hM4D(Gi)^+^ CNO^□^ p = 0.0001, hM4D(Gi)^+^ CNO^+^ VS hM4D(Gi)^□^ CNO^□^ p = 0.0001; error bars represent 95% confidence interval, CI). **f)** Cell-average mEPSC event frequency (One-way ANOVA, p = 0.3377). **g)** cumulative histogram of mEPSC amplitudes sampled from hM4D(Gi)^+^ CNO^+^ and hM4D(Gi)^□^ CNO^+^ conditions. Pink dashed line represents hM4D(Gi)^+^CNO^+^ distribution scaled according to the linear function *f*(*x*) = 0.6646*x* + 2.366. (Two-sample Kolmogorov-Smirnov test; Bonferroni correction α = 0.025; hM4D(Gi)^+^ CNO^+^ VS hM4D(Gi)^□^ CNO^+^, p = 6.2093e-07, Scaled VS hM4D(Gi)^□^ CNO^+^, p = 0.5231). **h)** biocytin fill of recorded pyramidal cell in GC co-expressing hM4D(Gi) and the GluA2-Ctail (Scalebar: 25µM). **i)** Top: representative mEPSC recordings; bottom: average mEPSC waveforms, black dashed line is aligned to GluA2-Ctail waveform peak. **j)** Cell-average mEPSC amplitudes (Empty Vector, n = 20; GluA2-Ctail, n = 17; Two-sample t-test, p = 0.0029). For electrophysiological data here and below, each point represents a recorded neuron and the mean and CI are represented by line and error bars.

### Neurons in the gustatory cortex express homeostatic synaptic scaling

The many hours-long time course over which CTA memory specificity emerges is reminiscent of the slow time course of homeostatic forms of plasticity such as synaptic scaling^11,12,28^. This suggests that synaptic scaling could serve as an important constraint on the Hebbian plasticity known to occur in the gustatory cortex following CTA^21,22^. Synaptic scaling has been extensively studied in the primary visual cortex and has been observed in other sensory cortices^15,29,30^, but whether neurons in the gustatory cortex are capable of expressing synaptic scaling is an open question. To test this, we took a chemogenetic approach using hM4D(Gi) DREADDS (designer receptors exclusively activated by designer drugs)^31^ to chronically inhibit pyramidal neurons in the GC and probe for the induction of synaptic scaling. Long-Evans rats received unilateral virus injections of AAV9-CAMKIIα-hM4D(Gi)-mCherry into the GC at p14 and after two weeks showed robust expression (Fig. 2a). In slice recordings from injected animals, acute application of CNO onto hM4D(Gi)^+^ neurons in GC resulted in hyperpolarization and a decrease in evoked spiking (Supp. Fig. 3), confirming the expression of hM4D(Gi). Using the protocol outlined in Figure 2b, animals were randomly assigned to the CNO control (CNO^□^) or CNO treatment (CNO^+^) groups. The contralateral, uninjected hemisphere of both CNO conditions served as an additional hM4D(Gi)^□^ control. After 2 days of chronic inhibition via CNO injection, we prepared brain slices from the GC and recorded miniature excitatory postsynaptic currents (mEPSCs) from mCherry^+^ pyramidal neurons to probe for the induction of synaptic scaling (Fig. 2c, Fig. 2d). We found a significant increase in mEPSC amplitude in the inhibited (hM4D(Gi)^+^ CNO^+^) neurons compared to both control groups (Fig. 2e) but no change in mEPSC frequency (Fig. 2f), as expected for classic synaptic scaling^11,13^. Analysis of the cumulative distribution function (CDF) of mEPSC amplitude from inhibited neurons revealed a significant shift towards higher amplitudes relative to control (hM4D(Gi)^□^ CNO^+^) (Fig. 2g). To test whether mEPSC amplitude increased multiplicatively, as is characteristic of synaptic scaling^11^, we plotted ranked inhibited vs. ranked control amplitudes and fit a linear function to the data^11,32^; scaling the inhibited distribution down generated a distribution that was statistically indistinguishable from the control (Fig. 2g). These results demonstrate that in response to activity perturbations, neurons in the gustatory cortex can homeostatically compensate through synaptic scaling.

**Fig. 3.**
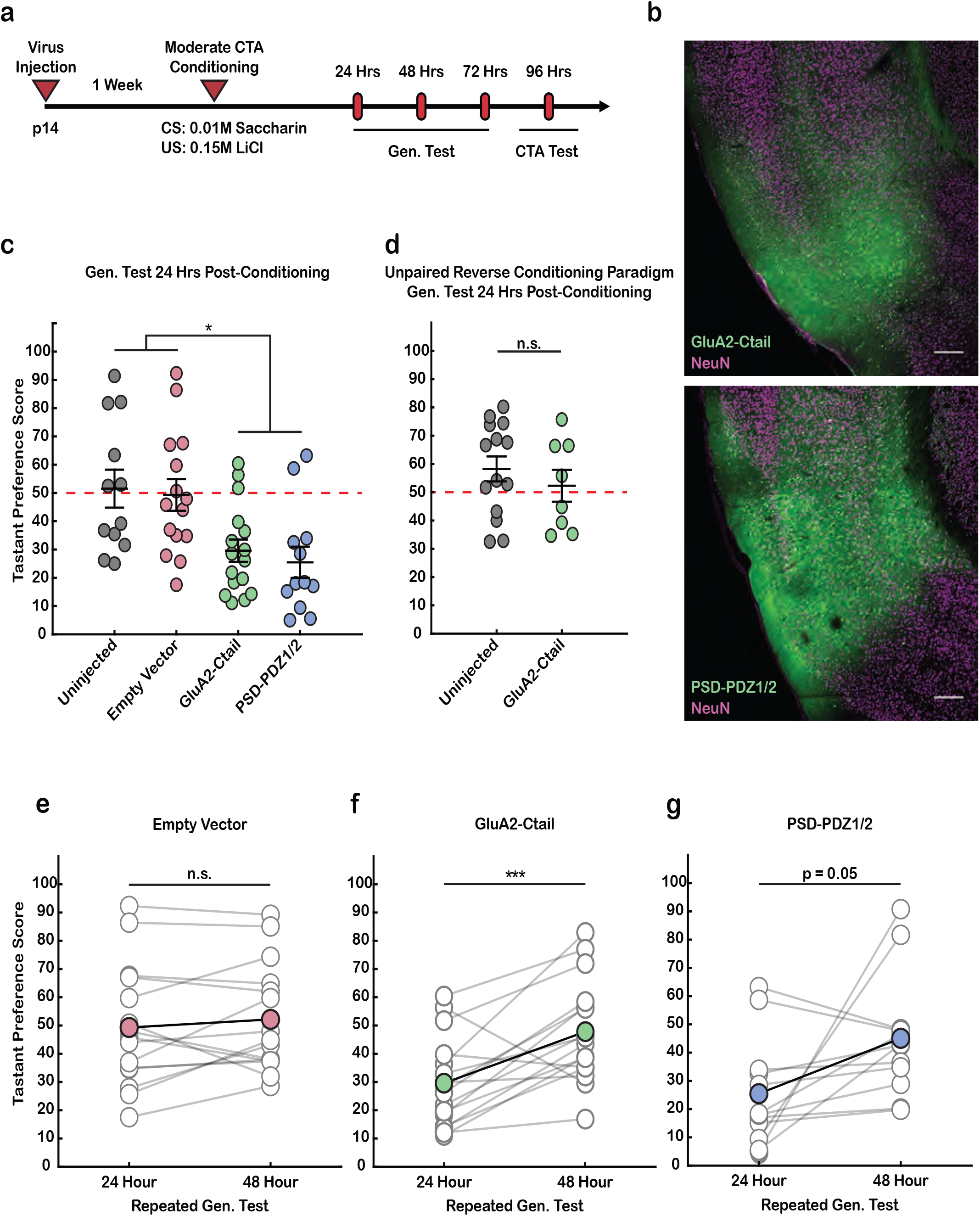
Perturbation of synaptic scaling extends CTA-induced generalized aversion. **a)** Basic experimental timeline. **b)** Top: Coronal brain slice depicting the gustatory cortex and expression of the GluA2-Ctail (Scalebar: 250 µM). Bottom: Coronal brain slice depicting the gustatory cortex and expression of the PSD-PDZ1/2 fragment (Scalebar: 250 µM). **c)** Gen. test, preference for NaCl tested 24Hrs following Moderate CTA induction (Uninjected, n = 12; Empty Vector, n = 15; GluA2-Ctail, n = 16; PSD-PDZ1/2, n = 12; One-Way ANOVA, p = 0.0015; Tukey-Kramer post-hoc test, UnInj. VS EV p = 0.9915, UnInj. VS Ctail p = 0.0289, UnInj. VS PSD p = 0.0124, EV VS Ctail p = 0.0401, EV VS PSD p = 0.0170, Ctail VS PSD p = 0.9490; error bars represent SEM). **d)** Gen. test, preference for NaCl tested 24Hrs following unpaired reverse conditioning paradigm (Uninjected, n = 14; GluA2-Ctail, n = 8; Two-sample t-test, p = 0.4206). **e-g)** Gen. test; preference for salt tested 24- & 48-hours post-conditioning (Empty Vector paired t-test, p = 0.3596; GluA2-Ctail paired t-test, p = 9.6572e-04; PSD-PDZ1/2 paired t-test, p = 0.0531)

Both synaptic scaling up and down are known to depend on C-terminal sequences on the GluA2 subunit of AMPA receptors^12,28,33,34,35^, and expression of the C-terminal fragment of GluA2 (the GluA2-Ctail) has been shown to block synaptic scaling in visual cortex pyramidal neurons^28,33^. To determine if synaptic scaling in GC pyramidal neurons is similarly dependent on GluA2 interactions, we next used a viral vector to express the GluA2-Ctail in GC. Rats received co-injections of AAV9-CAMKIIα-hM4D(Gi)-mCherry and either AAV2/1-GluA2-Ctail-GFP (GluA2-Ctail) or AAV2/1-GFP (Empty Vector) (Fig. 2h), were treated with CNO as above, and then recordings were obtained from hM4D(Gi)^+^ neurons ± the GluA2-Ctail (Fig. 2i). As expected, mEPSC amplitude was higher in inhibited neurons expressing the Empty Vector than in those expressing the GluA2-Ctail (Fig. 2j). Thus, synaptic scaling in GC relies on GluA2-Ctail interactions for its expression as it does in visual cortex^28,33^.

### Perturbation of synaptic scaling extends CTA-induced generalized aversion

We reasoned that if synaptic scaling shapes the specificity of CTA memory during the early stage of memory acquisition by constraining runaway LTP, then blocking synaptic scaling should prolong the expression of the generalized aversion. To test this hypothesis, we bilaterally expressed either the GluA2-Ctail or Empty Vector in the GC, subjected animals to moderate CTA conditioning and then probed for a generalized aversion (Fig. 3a,b top). As expected, 24 hours after conditioning the generalized aversion was gone in animals expressing Empty Vector, consistent with the result in our uninjected dataset (Fig. 1d and 3c); in contrast, in animals expressing the GluA2-Ctail, the generalized aversion was still evident at this late time point (Fig. 3c). Furthermore, this prolonged generalized aversion rapidly reversed after repeated exposure to the same tastant (Fig. 3f), similar to the extinction pattern observed at 24 hours post-conditioning in the Strong CTA group (Fig. 1g). However, the same testing paradigm induced no change in taste preference in the Empty Vector group, where the generalized aversion had already reversed (Fig. 3e). We then conducted an unpaired reverse conditioning paradigm in which animals received saccharin six hours after LiCl injection (Supp. Fig. 4a-c). In this paradigm, while animals were subject to both tastant experience and LiCl-induced malaise, no CTA was formed (Supp. Fig. 4d). GluA2-Ctail expression did not alter the preference to novel NaCl after reverse conditioning (Fig. 3d), confirming that the effect of the GluA2-Ctail relies on the induction of associative learning.

**Fig. 4.**
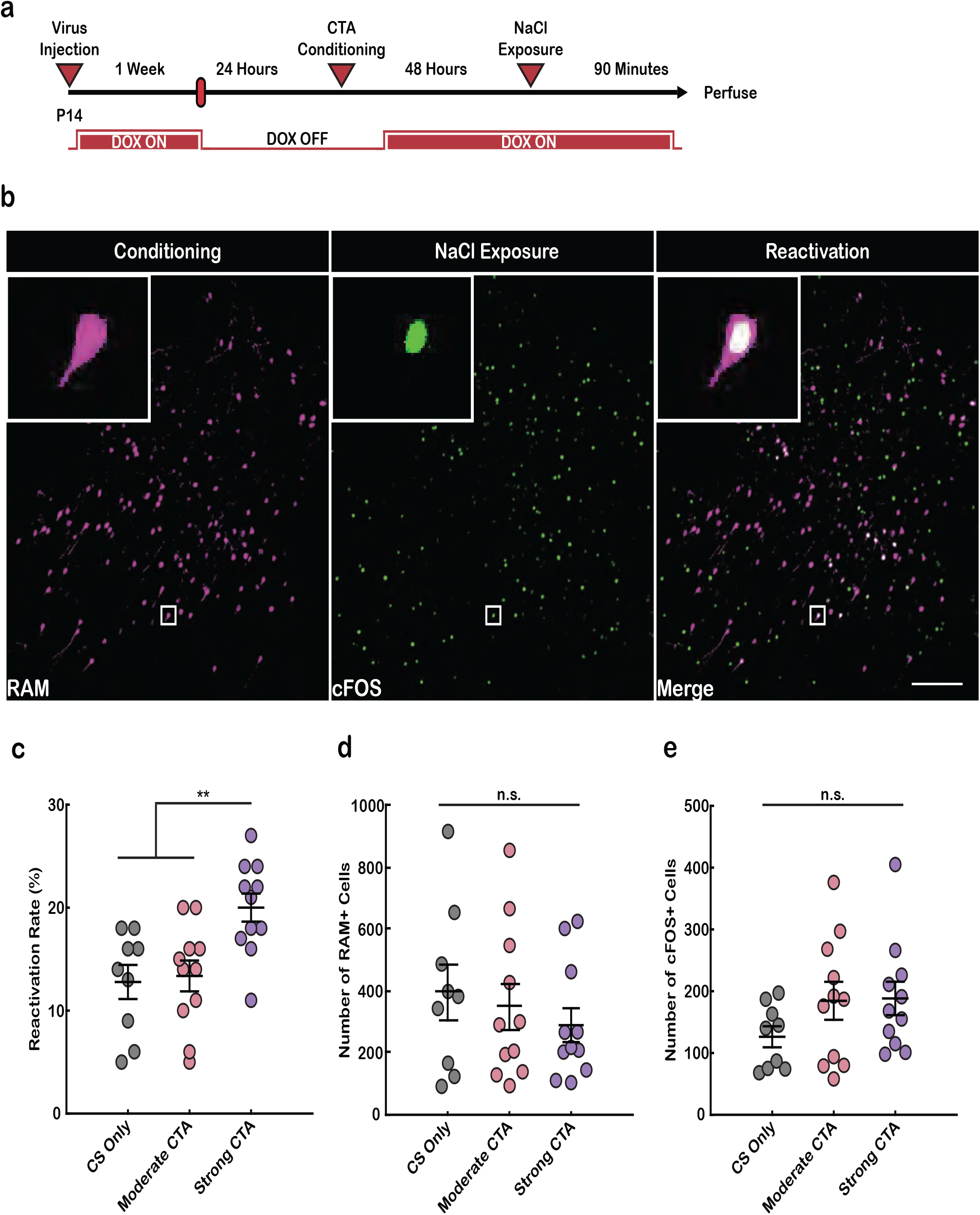
Conditioning-active GC neuronal ensemble are reactivated during generalized aversion. **a)** Experimental paradigm for labeling conditioning-active GC ensembles and their reactivation by the generalized tastant (NaCl). **b)** Representative images showing conditioning-active GC ensembles (RAM^+^, left), neurons activated by presentation of NaCl (c-FOS^+^, middle), and their overlap (RAM^+^c-FOS^+^, right; scalebar: 200 µM). **c)** Reactivation rate of conditioning-active neurons (RAM^+^c-FOS^+^/RAM^+^; CS Only, n = 9; MCTA, n = 11; SCTA, n = 11; One-Way ANOVA, p = 0.0027; Tukey-Kramer post-hoc testing, CS Only VS MCTA p = 0.9604, CS Only VS SCTA p = 0.0065. MCTA VS SCTA p = 0.0085; error bars represent SEM). **d)** Number of RAM^+^ cells (Kruskal-Wallis, p = 0.7598). **e)** Number of c-FOS^+^ cells (One-Way ANOVA, p = 0.2238).

Scaling up and down both depend on GluA2-Ctail interactions^12,28,33,34,35^. To differentiate between a role for scaling up and down in CTA specificity we next used a manipulation that specifically blocks scaling down, expression of the PDZ1/2 domains of PSD95 (PSD95-PDZ1/2)^36^. An AAV vector expressing PSD95-PDZ1/2 was bilaterally injected into GC (Fig. 3b bottom), and one week later we tested for the generalized aversion as described above. As with the GluA2-Ctail, PSD95-PDZ1/2 expression significantly prolonged the generalized aversion (Fig. 3c), and this reversed after repeated NaCl exposure (Fig. 3g). This finding demonstrates that disruption of synaptic scaling down in GC is sufficient to prevent the transition from a generalized to a conditioned stimulus-specific aversion.

Finally, as a third means of blocking synaptic scaling we used a GluA2 phosphorylation mutant (Y876E) that disrupts protein-protein interactions critical for synaptic scaling^37,38^. We bilaterally injected a lentiviral vector expressing either GluA2-Y876E or wild-type GluA2 (GluA2-WT) under control of a CAMKIIα promoter, and one week later subjected rats to our generalized aversion paradigm. While expression of GluA2-WT did not affect memory specificity, GluA2-Y876E expression prolonged the generalized aversion, which reversed after repeated NaCl exposure (Supp. Fig. 5a-c). None of these manipulations affected the consumption of saccharin during the conditioning trial, indicating these effects are not due to enhanced neophobia (Supp. Fig. 6a,b). In all cases, rats were left with an associative aversion to saccharin after generalized aversion testing was complete, demonstrating the consolidation of CTA memory (Supp. Fig. 6c,d). Because the CAMKIIα promoter used for the GluA2-Y876E construct mainly drives expression in cortical excitatory neurons^39^, these results suggest that homeostatic plasticity onto excitatory neurons in GC plays a crucial role in shaping stimulus specificity during CTA memory acquisition.

**Fig. 5.**
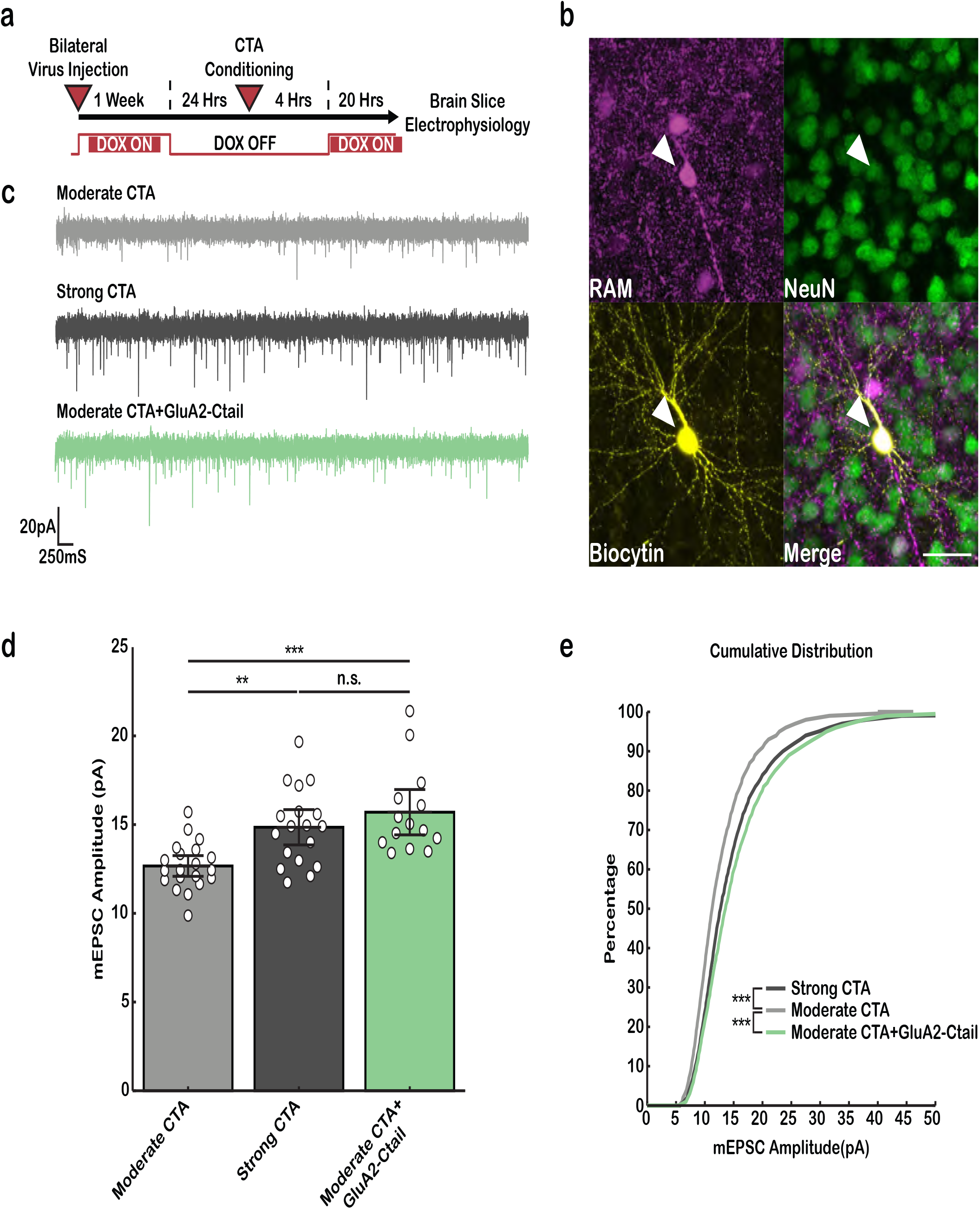
Blocking synaptic scaling prolongs the increase in synaptic strength onto CTA-active neurons following CTA-induction. **a)** Basic experimental timeline. **b)** Biocytin fill of pyramidal cell in GC expressing RAM (Scalebar: 25 µM) **c)** Representative mEPSC recordings. **d)** Cell-average mEPSC amplitudes of RAM^+^ neurons 24 hours after CTA conditioning (Moderate CTA, n = 20; Strong CTA, n = 18; Moderate CTA+GluA2-Ctail, n = 14; One-Way ANOVA, p = 1.0568e-04; Tukey-Kramer post-hoc test, Moderate CTA VS Strong CTA p = 0.0037, Moderate CTA VS Moderate CTA+GluA2-Ctail p = 0.0002, Strong CTA VS Moderate CTA+GluA2-Ctail p = 0.4516; error bars represent 95% CI) **e)** cumulative histogram of mEPSC amplitudes (Two-sample Kolmogorov-Smirnov test; Bonferroni correction α = 0.025; Moderate CTA VS Strong CTA p = 1.5833e-06, Moderate CTA VS Moderate CTA+GluA2-Ctail p = 1.0611e-11).

### Conditioning-active GC neuronal ensemble are reactivated during generalized aversion

A subset of neurons activated during conditioning will form the engram, a distributed memory trace that is reactivated during memory retrieval and is important for the expression of the memory^40^. The generalized aversion presumably occurs because – after conditioning – a novel tastant (such as NaCl) can activate these same ensembles that encode the aversive memory. We therefore wondered whether, during generalized aversion, ensembles of neurons within GC activated by NaCl might overlap more strongly with ensembles activated by conditioning. To label conditioning-activated neuronal ensembles and track their reactivation, we virally expressed a Robust Activity Marking (RAM) system in GC^41^. RAM consists of a tTA element driven by the synthetic activity-dependent promoter P_RAM_ and a TRE-dependent tdTomato marker; doxycycline prevents tTA from interacting with TRE, thereby inhibiting the expression of tdTomato. By removing doxycycline prior to conditioning and then restoring it afterwards, expression of tdTomato can be restricted to neurons active during conditioning.

We next took advantage of our ability to manipulate the duration of generalized aversion by controlling the intensity of CTA: generalized aversion induced by moderate conditioning is gone by 48 hours, while that induced by strong conditioning persists at this time point (Supp. Fig. 7) The longer interval between the conditioning trial and the generalization test ensures that doxycycline fully inhibits further expression of tdTomato. Animals first underwent moderate or strong conditioning, and 48 hours later were exposed to the novel tastant NaCl; endogenous expression of the immediate early gene c-FOS was used to label neurons activated by the NaCl exposure (Fig. 4a). By comparing the degree to which a novel tastant reactivates the conditioning-active ensemble during moderate and strong CTA training, we can examine reactivation in the absence and presence of the generalized aversion (Fig. 4b). Notably, we found that the reactivation rate of RAM-positive neurons during NaCl exposure was significantly higher in the Strong CTA than in the Moderate CTA or the CS Only groups (Fig. 4c). The distribution of RAM-positive neurons was similar among CS Only, Moderate CTA, and Strong CTA groups, consistent with previous reports that engram size remains constant despite changes in the strength of conditioning (Fig. 4d)^42,43^. Furthermore, the number of c-FOS-positive neurons in GC was not significantly different across experimental groups, indicating that the expression of generalized aversion is not determined by the size of active neuronal ensembles during NaCl exposure (Fig. 4e). We confirmed that the overlap between RAM-positive neurons and cFOS-positive neurons was above chance level in all groups (Supp. Fig. 8). In sum, our results demonstrate that during the expression of generalized aversion, a novel tastant induces robust reactivation of the conditioning-active GC ensemble. This result suggests that neuronal ensembles in GC that comprise the CTA engram may undergo changes in synaptic strengths that then allow non-conditioned tastants to activate them.

### Blocking synaptic scaling produces a persistent increase in synaptic strength following CTA conditioning

The duration of the generalized aversion is prolonged when synaptic scaling is blocked, and the conditioning-active neuronal ensemble in GC is reactivated by novel, unconditioned tastants during the generalized aversion. We hypothesized that the excitability of the conditioning-active ensemble is enhanced during CTA, and that this excitability enables the generalized aversion; subsequent homeostatic scaling down of synaptic strengths would then be expected to reduce excitability, thereby sculpting the specificity of CTA memory. This hypothesis predicts that synaptic strength onto conditioning-active ensembles should be potentiated when the generalized aversion is expressed and reduced again as the generalized aversion fades and memory specificity is established. To test this, we used RAM to label the conditioning-activated neuronal ensembles in GC (Fig. 5a), and then prepared brain slices and recorded from RAM^+^ (tdTomato^+^) neurons (Fig. 5b) 24 hours after Moderate or Strong CTA induction to compare synaptic strengths in the presence or absence of generalized aversion (Fig. 5c). We found that mEPSC amplitude was significantly higher in the Strong CTA condition compared to that of the Moderate CTA group (Fig. 5d). Moreover, mEPSC amplitudes onto RAM^+^ cells in the Moderate CTA group were similar to those from control, uninfected neurons (Fig. 2e).

Next, we asked whether excitatory synaptic strengths onto RAM^+^ neurons could be enhanced after Moderate CTA by blocking synaptic scaling using the GluA2-Ctail. Indeed, RAM^+^ neurons in the Moderate CTA+GluaA2-Ctail group had significantly higher mEPSC amplitudes than RAM^+^ neurons in the Moderate CTA condition alone and were not significantly different from the amplitudes observed in the Strong CTA condition (Fig. 5d). Analysis of the CDF demonstrated that, relative to the Moderate CTA group, both the Strong CTA and Moderate CTA+GluaA2-Ctail conditions were significantly shifted to the right, to higher amplitude values (Fig. 5e). These data suggest that synaptic strengths onto conditioning-active ensembles are normally potentiated and then homeostatically downscaled during Moderate CTA learning; blocking synaptic scaling in GC keeps synaptic strengths onto these conditioning-activated ensembles potentiated (Fig. 5d), and prolongs the generalized aversion measured behaviorally (Fig. 3).

## Discussion

Whether homeostatic synaptic scaling is capable of constraining Hebbian plasticity to enhance memory specificity has been a matter of much debate^44^. Here we took advantage of a paradigm (CTA) in which memory specificity develops slowly over many hours post-conditioning, to explore the mechanisms that reduce generalization and allow this specificity to emerge. We found that synaptic scaling is expressed by GC neurons, and that blocking synaptic scaling within GC prolongs memory generalization. Further, during generalized aversion, the novel tastant could reactivate CTA-active neuronal ensembles within GC, and this was correlated with enhanced synaptic strength onto these CTA-active ensembles. Finally, blocking synaptic scaling prolonged this enhancement of synaptic strengths onto CTA-active ensembles, in parallel with the behavioral prolongation of the generalized aversion. Taken together our data suggest that the enhancement of synaptic drive onto CTA-active ensembles (likely due to Hebbian plasticity^21,22^) initially enables a generalized aversion by allowing novel tastants to activate these ensembles; over time, synaptic scaling then downscales synapses onto these ensembles and this process establishes the specificity of the CTA memory. Our data also show that the duration of generalization aversion depends on both the strength of the CTA induction, and the magnitude of synaptic scaling. This raises the interesting possibility that the relative strengths of Hebbian and homeostatic plasticity can be tuned to control the degree to which an associative memory can be generalized.

Extensive theoretical work has explored the idea that Hebbian forms of plasticity (such as LTP) can destabilize neural networks by saturating synaptic weights, and that the addition of synaptic scaling can counteract this instability to help establish or maintain memory specificity^8,9,10,17^. There is some experimental evidence that blocking pathways important for synaptic scaling can influence memory^45,46^, but it was not clear from these earlier studies whether these manipulations impacted the transition from generalized to specific memories following the induction of learning. A major advantage of the paradigm we establish here is that initial formation of CTA memory is rapid and then followed by a slower process that establishes the specificity of the memory, allowing us to dissociate these two processes. We found that manipulations that block synaptic scaling had no impact on initial CTA memory formation, which depends on NMDAR-dependent plasticity^21^; this is consistent with the slow time course of synaptic scaling, which theoretical work suggests should not be able to fully constrain Hebbian plasticity in the short term^18^. Rather than limiting initial memory formation, our data show that synaptic scaling is important for controlling the slow transition from generalized to specific memories. It is possible that homeostatic synaptic scaling in cortical networks also contributes to other aspects of memory encoding, such as memory consolidation, that unfold over slower timescales^47^.

Synaptic scaling has been extensively studied in sensory cortices where is it straightforward to induce it through manipulations of sensory input^5,29,30^, but has not previously been demonstrated in other neocortical areas. To probe for synaptic scaling in GC neurons we used viral expression of the hM4D(Gi) DREADD in GC to chronically inhibit excitatory neurons. This paradigm induced robust multiplicative synaptic scaling that was dependent on GluA2-Ctail interactions, indicating that it shares key molecular features with synaptic scaling in other neocortical areas^28,33^. We also found that excitatory synapses onto GC neurons active during CTA induction were potentiated in experimental conditions associated with the expression of a generalized aversion; this potentiation was gone when CTA memory became specific for the conditioned tastant, and was prolonged in CTA-active neurons expressing synaptic scaling blockers. Taken with the ability of these blockers to prolong the behaviorally measured generalized aversion, this strongly suggests that synaptic scaling is the critical cellular mechanism in GC neurons that modulates the specificity of CTA memory. However, there is overlap in the molecular machinery that supports synaptic scaling and other forms of synaptic depression. Some forms of LTD also depend on GluA2 C-tail interactions^48,49^, while the role of GluA2 Y876 phosphorylation is less clear and likely depends on the experimental paradigm used^49,50,51,52^. Behaviorally, expression of the GluA2-Ctail can block memory processes thought to depend on LTD, including forgetting^53,54^ and extinction^55^, while sparing memory formation^56,57^. In contrast, LTD and scaling down rely on distinct domains of PSD95: the N-terminal PDZ1/2 domains of PSD95 are required for synaptic scaling down^36,58^, while its C-terminal Src homology 3(SH3) and guanylate kinase (GK) domains are critical for induction of LTD^59,60^. The common feature of all three manipulations we used to extend the CTA-induced generalized aversion is their ability to block synaptic scaling down; taken together with our physiological demonstration of synaptic scaling down, these data strongly suggests that memory specificity is established through synaptic scaling down, rather than LTD. Furthermore, overexpression of scaling blockers did not impair acquisition of CTA memory, which depends on NMDAR-dependent plasticity^21^. Our data thus support the hypothesis that memory formation is initiated and established through Hebbian mechanisms, and its specificity is then fined-tuned by synaptic scaling.

The prevailing view of associative memory formation is that a subset of neurons activated during conditioning will undergo enduring changes and become a memory engram^40^. Apart from enabling memory retrieval, subpopulations of these conditioning-active ensembles may also regulate the generalizability of memory^61^. In our CTA paradigm, we found that the conditioning-active GC neurons are robustly reactivated by a novel (generalized) tastant only during behavioral states in which the animals exhibit a generalized aversion. The degree of reactivation by the generalized tastant is comparable to the rate of reactivation by specific memory recall observed in other learning paradigms^62^. More importantly, we showed that excitatory postsynaptic strength onto these conditioning-active neurons directly correlated with the expression of generalized aversion; preventing synaptic scaling down specifically in GC increased the duration of this postsynaptic enhancement, and also prolonged generalization. Taken together, this suggests that conditioning-active GC ensembles are a critical neuronal population that undergo plastic changes that contribute to the expression and decay of generalized aversion.

While it has long been known that CTA can generalize to other novel tastants^24,25^, the cellular mechanism that control the specificity of CTA memory have remained obscure. Here we provide an important advancement by demonstrating that the duration of the generalized aversion is controlled by a homeostatic process that renormalizes synaptic strengths onto conditioning-active cortical ensembles to establishes the tastant specificity of CTA. Our data are consistent with a model in which unopposed Hebbian plasticity^21,22^ onto GC neurons is important for initial CTA memory formation and generalization, while subsequent homeostatic synaptic scaling down slowly restores excitability and sculpts memory specificity. The initial generalization of CTA enabled by slow homeostatic compensation might be ethologically useful, by encouraging caution toward novel foods in an environment where such foods have recently proven dangerous. Conversely, if left unchecked persistent generalization of aversive conditioning could become pathological, as in post-traumatic stress disorders^63^. Our data suggest that the degree of specificity of CTA memory is malleable and can be controlled by the relative strengths of Hebbian and homeostatic plasticity within GC.

## Acknowledgments

We thank Edwin Zhang for assistance with behavioral experiments and histology, and Brian Cary for help with data analysis. We thank members of both the Turrigiano and Katz lab for their feedback on this work. Supported by NIH grants 1 F31 NS108506-01 (RR), R35NS111562 (GGT), and R01DC006666 (DBK).

## Author Contributions

Conceptualization, C.H.W., R.R., D.B.K., & G.G.T.; Methodology, C.H.W., R.R., D.B.K., & G.G.T.; Investigation, C.H.W. & R.R.; Formal Analysis, C.H.W. & R.R.; Writing – Original Draft, C.H.W., R.R., & G.G.T.; Writing – Review and Editing, C.H.W., R.R., D.B.K., & G.G.T.; Funding Acquisition, R.R., D.B.K., & G.G.T.; Resources, D.B.K., & G.G.T.; Supervision, D.B.K., & G.G.T.

## Methods

All experimental procedures were approved by Brandeis University Institutional University Animal Care and Use Committee and followed the National Institute of Health guideline for the Care and Use of Laboratory Animals.

## Rats

Young Long-Evans rats (p28-p34) were used in these experiments. Timed pregnant rats were obtained from Charles River Laboratories, and the progeny were maintained in Foster Biomedical Research Labs at Brandeis University. After weaning at post-natal day 21 (p21), littermates were individually housed in a humidity- and temperature-controlled environment and entrained to a 12 hour light–dark cycle (light phase from 7:00-7:00) with ad libitum access to food and water unless described otherwise. In all behavioral experiments, because there were no sex differences in taste preference during Gen. test (24 hours post-conditioning), rats of both sexes were randomly assigned to different experimental conditions. For experiments that required virus-mediated manipulations, virus surgery was performed on animals at p14, which allowed the construct to be fully expressed at the time of conditioning (∼p28). The average infection rates for neurons in GC for GluA2-Ctail and PSD-PDZ1/2 were 35.5% and 26% respectively (n = 3 for each condition), calculated as GFP^+^NeuN^+^ cells / NeuN^+^ cells (%). For all behavioral experiments, viral expression was confirmed post hoc by immunostaining and all animals with detectable bilateral expression in GC were included for analysis. All subjects selected for electrophysiology experiments were age matched to animals selected for behavioral experiments.

## Viral vectors

The pAAV-CMV-GluA2-CT (GluA2-Ctail) and pAAV-CMV-PSD95-PDZ1/2 (PSD-PDZ1/2) plasmids were constructed by sub-cloning the coding sequences of the scaling blockers^32,35^ into the pAAV-CMV-eGFP3 vector (Empty Vector). pAAV-RAM-dtTA-TRE-tdTomato was constructed by replacing GFP in pAAV-RAM-dtTA-TRE-GFP (Addgene: 84469) with the coding sequence of tdTomato. pAAV-CAMKIIα-hM4D(Gi)-mCherry was obtained from Addgene (171074). Lenti-CAMKIIa-GFP-GluA2-Y876E and Lenti-CAMKIIa-GFP-GluA2-WT were generated by subcloning coding sequences of full-length GluA2 into a pLenti-CAMKIIa-c1v1 vector. For *in vivo* applications, GluA2-Ctail, PSD-PDZ1/2, and Empty Vector were packaged in AAV2 serotype 1; RAM-dtTA::TRE-tdTomato and hM4D(Gi)-mCherry were packaged in AAV2 serotype 9. All viruses were produced at Duke Viral Vector Core, except for the GluA2-Ctail, which was packaged at UPenn Viral Vector Core.

## Virus Surgery

Rats were anesthetized with a cocktail containing ketamine (70 mg/kg, intraperitoneally (i.p.)), xylazine hydrochloride (3.5 mg/kg), and acepromazine maleate (0.7 mg/kg), and placed onto a stereotaxic apparatus. The skull was exposed, and craniotomies were made above the GC. For the behavior experiments (GluA2-Ctail, PSD-PDZ1/2, Empty Vector) and imaging (RAM), viruses (800 nl per hemisphere) were bilaterally microinjected into the GC through a glass micropipette connected to a micromanipulator (Narishige, MO-10), at a rate of approximately 200 nl/min. To cover the whole GC, three injection sites were chosen for each hemisphere: anterior-posterior (AP) with reference to bregma: 1.0 mm, medial-lateral (ML): ±4.7 mm, dorsal-ventral (DV) with reference to the brain surface: -3.5 mm, -3.6 mm, -3.7 mm. To allow adequate diffusion of virus particles, the pipet remained in place for additional 5 minutes after injection and was slowly withdrawn from the site. For DREADDS experiments, hM4D(Gi)-mCherry (400 nl) was unilaterally injected into GC. To label conditioning-active neurons for electrophysiological experiments, either RAM alone (400 nl), or a cocktail containing RAM and GluA2-Ctail (1:1, 400 nl) were bilaterally infused.

## Behavioral paradigm

### Two-Bottle Paradigm

This CTA behavioral paradigm was adapted from Flores et al., 2016 and modified for our experimental needs. After being transferred into individual home cages, rats were habituated to two bottles with *ad libitum* access to water for three days. The animals were then subjected to water restriction for an additional three days, during which the access to water was limited to two hours. On the fourth day of restriction, rats underwent CTA conditioning. They received two bottles that contained the conditioned stimulus (CS), saccharin (10 mM), for thirty-minutes, followed by an intraperitoneal injection of the unconditioned stimulus (US), LiCl (for moderate conditioning, 0.15 M; for strong conditioning, 0.30 M, 1% Body Weight). For the CS Only group, rats received saccharin, and were injected with saline instead of LiCl. For the US Only groups, rats were given two bottles of water during the conditioning trial, followed by an injection of LiCl. After the conditioning, rats underwent a retention interval of 4 or 24 hours until testing. For a two-bottle choice test, rats were given one bottle of tastant, counterbalanced by one bottle of water, for thirty minutes. The results were quantified using a tastant preference score (TPS):

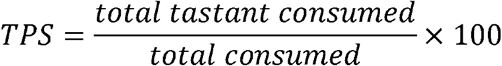

To test CTA (CTA test), saccharin (10 mM) was used as the tastant. To test generalized aversion (Gen. test), NaCl (150 mM) was used. To measure the attenuation and reversal of the generalized aversion, Gen. tests were conducted daily for three days. After Gen. testing was complete, rats were given a CTA test to ensure that the animals had indeed learned an aversion to the CS. All the consumption was documented throughout the paradigm to ensure that daily fluid intake was stable.

### Reverse conditioning

The same two-bottle training paradigm was conducted with the following modification: on the day of conditioning, rats first received injection of LiCl (0.15 M), and six hours later, 30-minute access to saccharin. Gen tests were then conducted the next day to match the 24-hour retention interval.

### Administration of clozapine N-oxide (CNO)

Rats underwent surgery as described above and were individually housed at p21. At p27-p30, CNO (3 mg/kg) was intraperitoneally administered every 12 hours for 2 days. Animals assigned to the DREADDS-only group underwent virus injection, but CNO was replaced with saline during the drug administration. After the treatment, acute brain slices were collected for electrophysiological analysis.

### Labeling of conditioning-active neurons

Customized chow containing low-dose doxycycline (40 ppm, ScottPharma) were added to the home cage one day before virus surgery, and rats were maintained on doxycycline throughout the training. The doxycycline-containing chow was replaced with regular chow one day before the conditioning trial to allow adequate RAM induction. Two hours after the conditioning trial, rats were placed back on a diet containing high-dose doxycycline (100 ppm, ScottPHarma) to prevent further RAM activation. To test the reactivation of conditioning-active ensembles, rats underwent the training paradigm as described above, except that they were given NaCl for thirty minutes 48 hours after the conditioning trial. Ninety minutes after the exposure, the animals were sacrificed for further immunohistochemistry experiments. For the electrophysiological experiments, acute brain slices were collected 24 hours after the conditioning trial.

## Immunohistochemistry

Animals were deeply anesthetized with isofluorane and perfused with 4% paraformaldehyde (PFA). Brains were extracted, post-fixed in 4% PFA for 48 hours, and then sliced on a vibrating microtome (Leica Vibratome VT 1200s). Coronal brain slices (50 µm) containing GC were collected serially and stored in PBS until staining. For immunostaining, 2 slices were selected from anterior, middle, and posterior GC (6 slices in total) from each animal. Floating slices were washed three times with PBS, preincubated with blocking buffer (5% goat serum/3% BSA/0.3% Triton X-100 in PBS) at room temperature for 2 hours, and then incubated with primary antibodies diluted in blocking buffer at 4°C overnight. To verify expression of scaling blockers, chicken anti-GFP (1:1000, Aves Labs) and mouse anti-NeuN (1:500, MAB-377, Millipore) were used. To verify expression of hM4D(Gi), we used rat anti-mCherry (1:1000, Thermo-Fisher) and mouse anti-NeuN (1:500, MAB-377, Millipore). To label the reactivation of conditioning-active ensembles, mouse anti-RFP (1:1000, Rockland) and rabbit anti-c-FOS (1:200, 9F6, Cell Signaling Technology) were used. On the next day, slices were first washed thoroughly with PBS 5 times, and then incubated with Alexa-conjugated secondary antibodies diluted in PBS containing 5% goat serum and 3% BSA at room temperature for 3 hours (goat anti-chicken Alexa-488, goat anti-mouse Alexa-594, goat anti-rat Alexa-594, goat anti-mouse Alexa-555, and goat anti-rabbit Alexa -647, 1:400, Thermo-Fisher). After 3 more washes with PBS, slices were either directly mounted onto the slides and cover slipped using DAPI-Fluoromount-G mounting medium (SouthernBiotech), or counterstained with Hoechst stain (1:1000, Thermo-Fisher) for 20 minutes before mounting with Fluoromount-G medium (SouthernBiotech).

## Image acquisition and analysis

Images were acquired on a laser-scanning confocal microscope (Zeiss LSM880) using ZEN Black acquisition software (Zeiss). The boundaries of GC were manually determined based on the Paxinos and Watson rat brain atlas (Paxinos & Watson, 2013). Images were obtained using tile scan under a 20x objective, with frame size of 512 × 512. For all experiments, acquisition settings including laser power, gain/offset and pinhole size were kept consistent. To quantify reactivation of conditioning-active ensembles, image tiles were first subjected to maximum intensity projection and stitch functions using ZEN Black, then analyzed using ImageJ software (NIH, US). Images from each channel were background-subtracted using the rolling ball function, and then thresholded to outline RAM^+^, c-FOS^+^, and DAPI^+^ cells. The rostral-to-caudal distribution of RAM^+^ neurons varied slightly due to the efficacy of virus spread. Therefore, for each animal, 3 consecutive hemispheres that showed highest RAM expression were included for quantification. ROI of the same size (2 mm^2^) was drawn to include GC for all hemispheres across experimental groups; the average numbers of RAM^+^, c-FOS^+^, and DAPI^+^ cells from all hemispheres were quantified using particle analysis function, and the reactivation rate was calculated as follows:

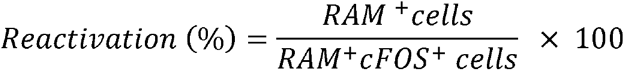

The number of double-labeled cells compared with the chance level was also calculated to ensure that the reactivation was not due to random overlap (Khalaf et al., 2018):

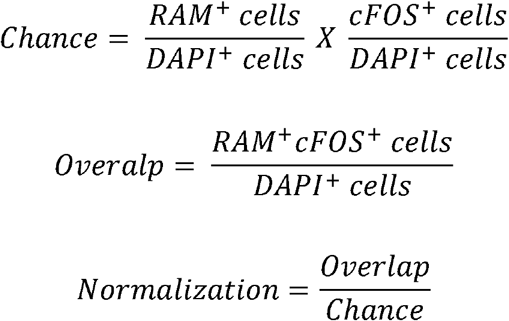

## Electrophysiology

### Standard Protocols

#### *Ex-Vivo* Acute Brain-Slice Preparation

Brain slices were produced following our previously documented protocols (Miska et al., 2018; Cary & Turrigiano, 2019). Briefly, rats (p28-p32) were anesthetized with isoflurane, decapitated, and the brain was swiftly dissected out in ice cold carbogenated (95% O2, 5% CO2) standard ACSF (in mM: 126 NaCl, 25 NaHCO3, 3 KCl, 2 CaCl2, 2 MgSO4, 1 NaH2PO4, 0.5 Na-Ascorbate, osmolarity adjusted to 310-315 mOsm with dextrose, pH 7.35). Coronal brain slices (300 μm) containing the gustatory cortex were obtained from both hemispheres of each animal using a vibratome (Leica VT1000). The slices were immediately transferred to a warm (34°C) chamber filled with a continuously carbogenated ‘protective recovery’ (Ting et al., 2014) choline-based solution (in mM: 110 Choline-Cl, 25 NaHCO3, 11.6 Na-Ascorbate, 7 MgCl2, 3.1 Na-Pyruvate, 2.5 KCl, 1.25 NaH2PO4, and 0.5 CaCl2, osmolarity 310-315 mOsm, pH 7.35) for 10 min, then transferred back to warm (34°C) carbogenated standard ACSF and incubated another 45 min. Brain slices were used for electrophysiology experiments between 1 – 7 Hours post-slicing.

#### Whole-Cell Recording

Slices were visualized on an Olympus upright epifluorescence microscope using a 10x air (0.13 numerical aperture) and 40x water-immersion objective (0.8 numerical aperture) with infrared-differential interference contrast optics and an infrared CCD camera. Gustatory cortex was identified in acute slices using the shape and morphology of the corpus callosum, piriform cortex and the lateral ventricle as a reference. The borders of GC were determined by comparing the aforementioned landmarks in slice to the Paxinos and Watson rat brain atlas. Pyramidal neurons from superficial and deep layers were visually targeted and identified by the presence of an apical dendrite and teardrop shaped soma. In experiments involving the expression of a viral construct, fluorophore expression was used to visually target pyramidal neurons. Morphology was further confirmed by post-hoc reconstruction of biocytin fills. Borosilicate glass recording pipettes were pulled using a Sutter P-97 micropipette puller, with acceptable tip resistances ranging from 3 to 6 MΩ. Inclusion criteria for neurons included V_m_, R_in_, and R_s_ cut-offs as appropriate for experiment type and internal solution. All recordings were performed on submerged slices, continuously perfused with carbogenated 35°C recording solution. Data were low-pass filtered at 10 kHz and acquired at 10 kHz with Axopatch 700B amplifiers and CV-7B headstages (Molecular Devices, Sunnyvale CA). Data were acquired using WaveSurfer v0.953 (Janelia Research Campus), and all post-hoc data analysis was performed using inhouse scripts written in MATLAB (Mathworks, Natick MA).

#### mEPSC recordings

When recording mEPSCs, Cs+ Methanesulfonate-based internal recording solution was used. This Cs+ internal was modified from Xue et al., 2014, and contained (in mM) 115 Cs-Methanesulfonate, 10 HEPES, 10 BAPTA·4Cs, 5.37 Biocytin, 2 QX314 Cl, 1.5 MgCl2, 1 EGTA, 10 Na2-Phosphocreatine, 4 ATP-Mg, and 0.3 GTP-Na, with sucrose added to bring osmolarity to 295 mOsm, and CsOH added to bring pH to 7.35. For these recordings, pyramidal neurons were voltage clamped to −70 mV in standard ACSF containing a drug cocktail of TTX (0.2 μM), APV (50 μM), PTX (25 μM). Traces of 10 seconds were acquired over a period of ∼10-15 minutes allowing for the cell to fill for later morphological verification. Neurons were excluded from analysis if Rs > 25 MΩ.

#### mEPSC analysis

To reliably detect mEPSC events and limit selection bias, we used in-house software that employs a semi-automated template-based detection method (Miska et al, 2018; Cary & Turrigiano, 2019). Event inclusion criteria included amplitudes greater than 5 pA and rise times less than 3 ms. The resulting events detected by our software were visually assessed for inclusion/exclusion. Our average manual exclusion rate across all experiments was ∼9% of events detected. Additionally, the automated detection method missed, on average, ∼6% of events, that were manually included. The experimenter was blinded to experimental condition and treatment until after the analysis was complete.

#### I-Clamp recordings

For I-clamp recordings, a K-gluconate internal recording solution was used. This internal contained (in mM) 100 K-gluconate, 10 KCl, 10 HEPES, 5.37 biocytin, 10 Na2-phosphocreatine, 4 Mg-ATP, and 0.3 Na-GTP, with sucrose added to bring osmolarity to 295 mOsm and KOH added to bring pH to 7.35. Pyramidal neurons in superficial and deep layers of gustatory cortex expressing inhibitory DREADDS + mCherry were targeted for whole cell recording. The slices were continuously perfused with standard ACSF containing a drug cocktail of APV (50 μM), PTX (25 μM), and DNQX (25 μM). A series of 20 5s traces were recorded; on odd numbered traces input resistance was assessed using a -100 pA, 500 ms DC current injection, while on even numbered traces DC current steps of varying amplitude (−60 to 300 pA) were given to asses firing rate as a function of injected current (FI curves). The V_r_ and FI curves of hM4D(Gi)^+^ neurons were assessed before & after perfusion of the exogenous DREADD agonist, CNO (1 μM).

#### I-Clamp analysis

Changes in input resistance pre and post CNO treatment were calculated using Ohm’s Law. Changes in V_r_ were quantified by analyzing the average resting membrane potential during the 1^st^ minute and 10^th^ minute of the CNO wash. Lastly, action potentials were detected using a custom Matlab script. The results of the spike detection function were visually assessed. Neurons were excluded if Rs > 25 MΩ or V_r_ > -50 mV.

#### Biocytin Reconstruction

After recording, slices were incubated in cold 4% PFA for two days. Following fixation slices were stained as described above. Biocytin fills were recovered by counterstaining with AlexaFluor streptavidin (ThermoFisher). Images were acquired using the Leica SP5 Laser Scanning Confocal Microscope.

#### Specifics regarding the chemogenetic induction, and blocking, of synaptic scaling

Acute brain slices were prepared as described above, with the exception that TTX was included in the standard ACSF used for slicing and incubation. This was done to prevent any plasticity induction that might occur during release from CNO inhibition in slices rendered hyperexcitable due to chronic inhibition. Pyramidal neurons in superficial and deep layers of gustatory cortex expressing mCherry were targeted for whole-cell patching and mEPSC recording as described above. For experiments involving the co-injection of hM4D(Gi) and the GluA2-Ctail, cells targeted for recording were confirmed to be expressing both mCherry and GFP post-hoc through immunostaining of cells using antibodies described in immunohistochemistry section.

#### Specifics regarding recordings from the conditioning-active ensemble

Slices were created exactly 24-hours post conditioning using the methods described above. mEPSCs were recorded using the method described above. For recording, fluorescent RAM^+^ (tdTomato^+^) cells were targeted in both superficial and deep layers, where expression was equally robust. For experiments involving the co-injection of RAM and the GluA2-Ctail, cells targeted for recording were confirmed to be expressing both tdTomato and GFP post-hoc through immunostaining of the cells using antibodies described in the immunohistochemistry section.

### Statistical analysis

For all experiments including behavior, electrophysiology and imaging, individual experimental distributions were tested for normality using the Anderson-Darling test. If all experimental conditions passed the normality test, a t-test, paired t-test, or one-way ANOVA were used where appropriate. Significant ANOVA tests were followed by Tukey-Kramer post hoc comparisons. If one or more conditions failed to pass the normality test, a Wilcoxon ranksum or Kruskal-Wallis were used as appropriate. Significant Kruskal-Wallis tests were followed by Bonferroni post hoc comparisons. Differences between cumulative distributions were tested using a two-sample Kolmogorov-Smirnov corrected for multiple comparisons. Results of all statistical tests can be found in the figure legends. For behavior experiments n = number of animals, while for electrophysiology experiments n = number of cells; these values are given in the figure legends. Electrophysiogical data were collected from at least 4 animals for each condition.

## Data availability

All data generated in this study are included in the figures and supplemental figures.

## Code availability

Custom MATLAB codes used for mEPSC analyses can be found at https://github.com/BrianAndCary/papers/tree/master/bcary2020_paper/mini_FI_GUI

