## Supplementary figures and images for "Homeostatic Synaptic Scaling Establishes the Specificity of an Associative Memory"

### Supplemental Figures

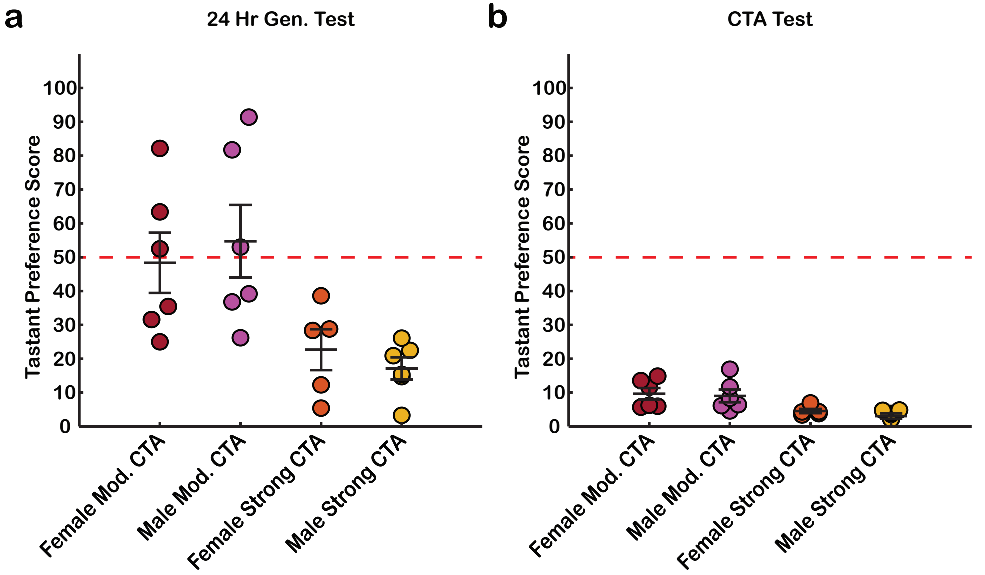

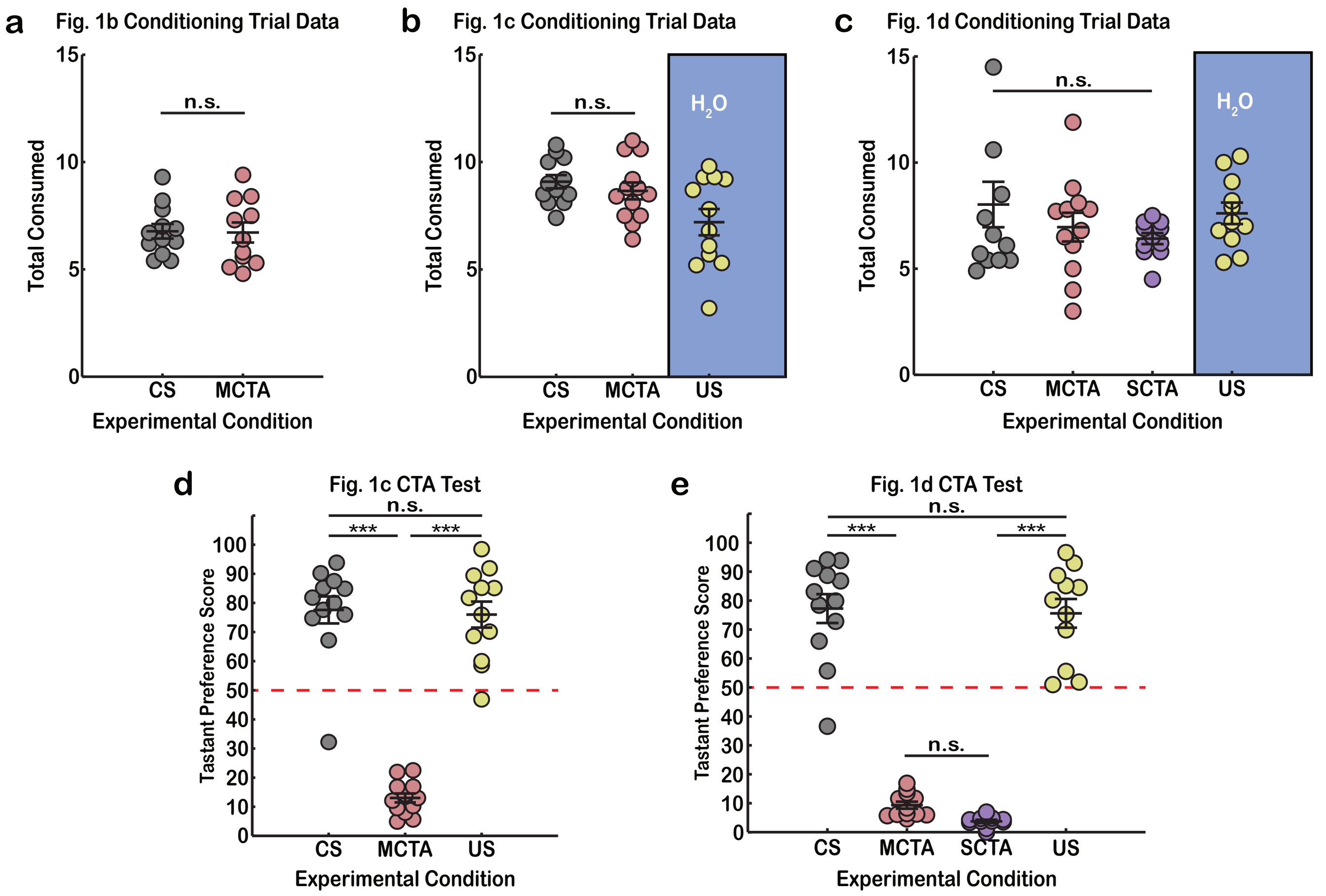

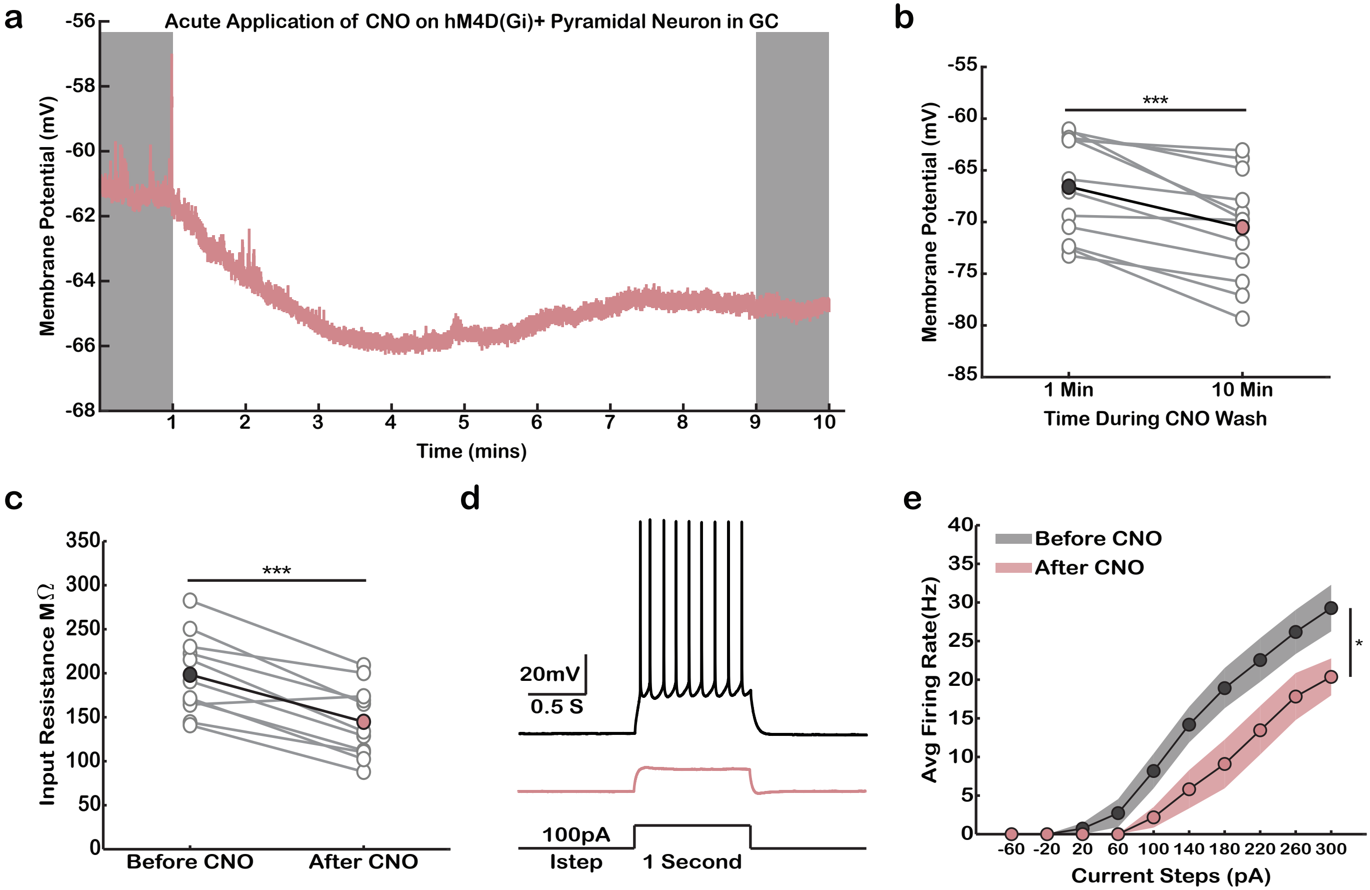

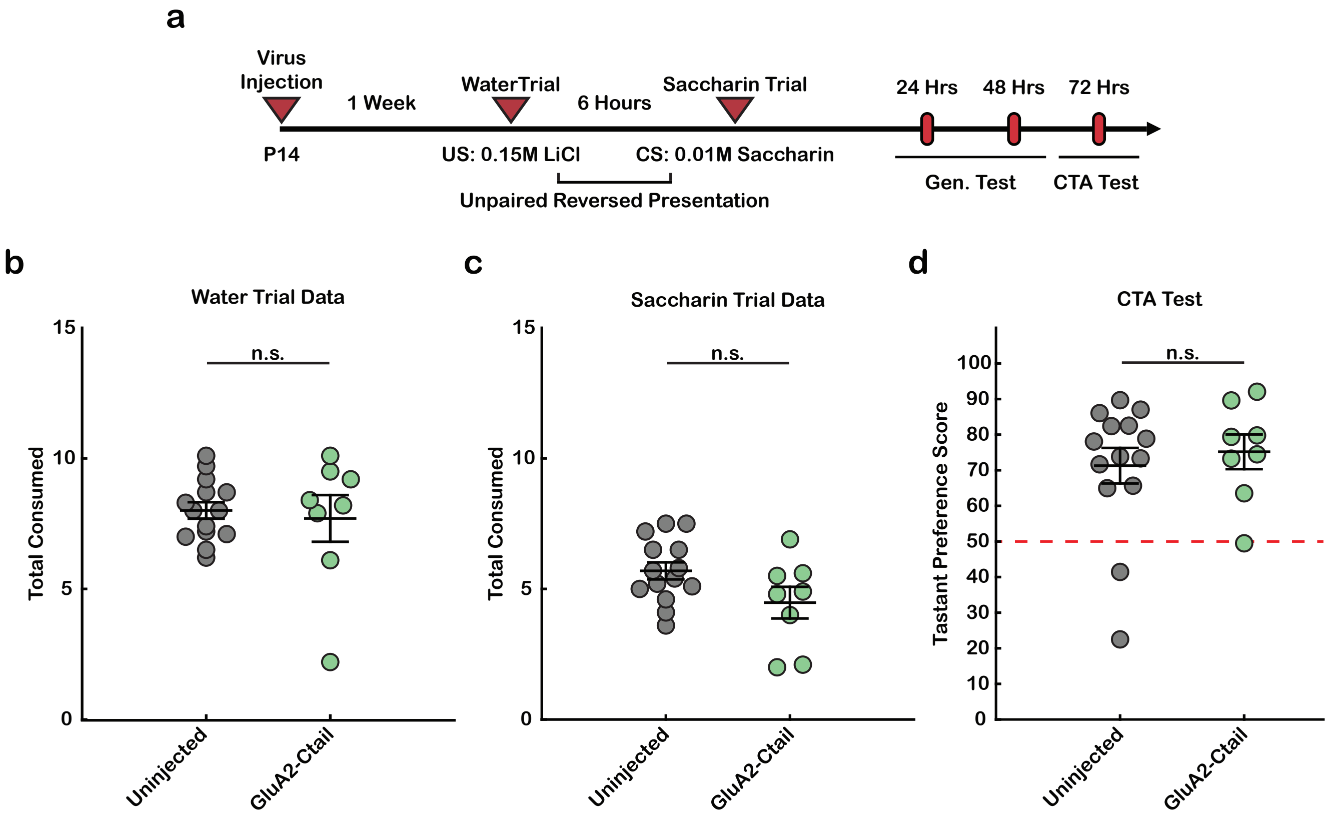

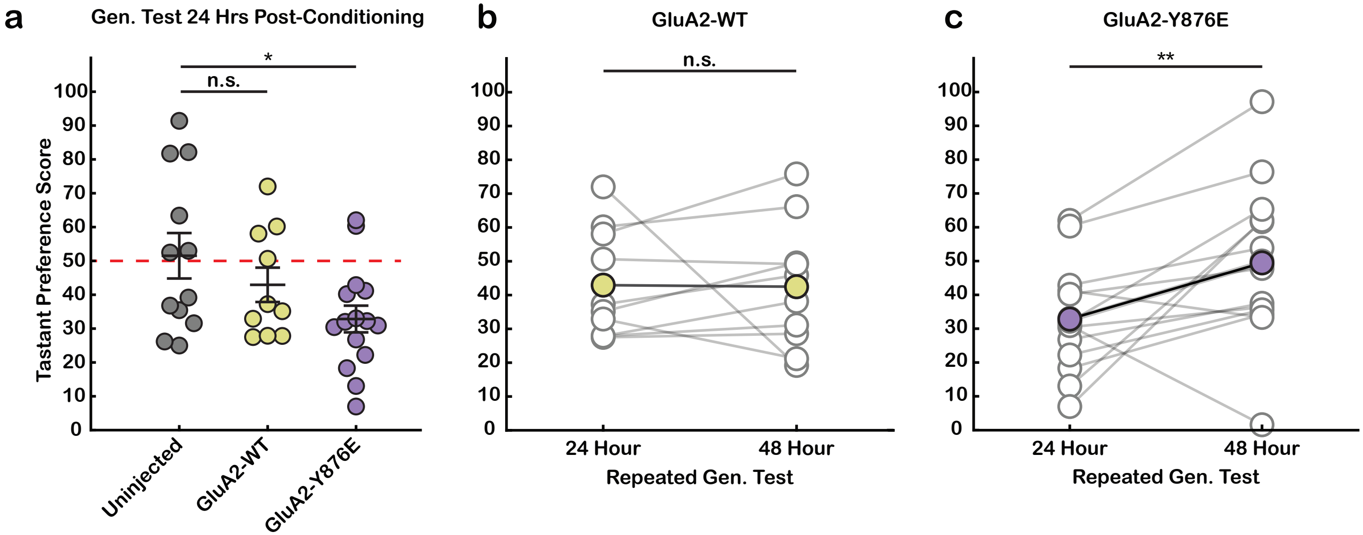

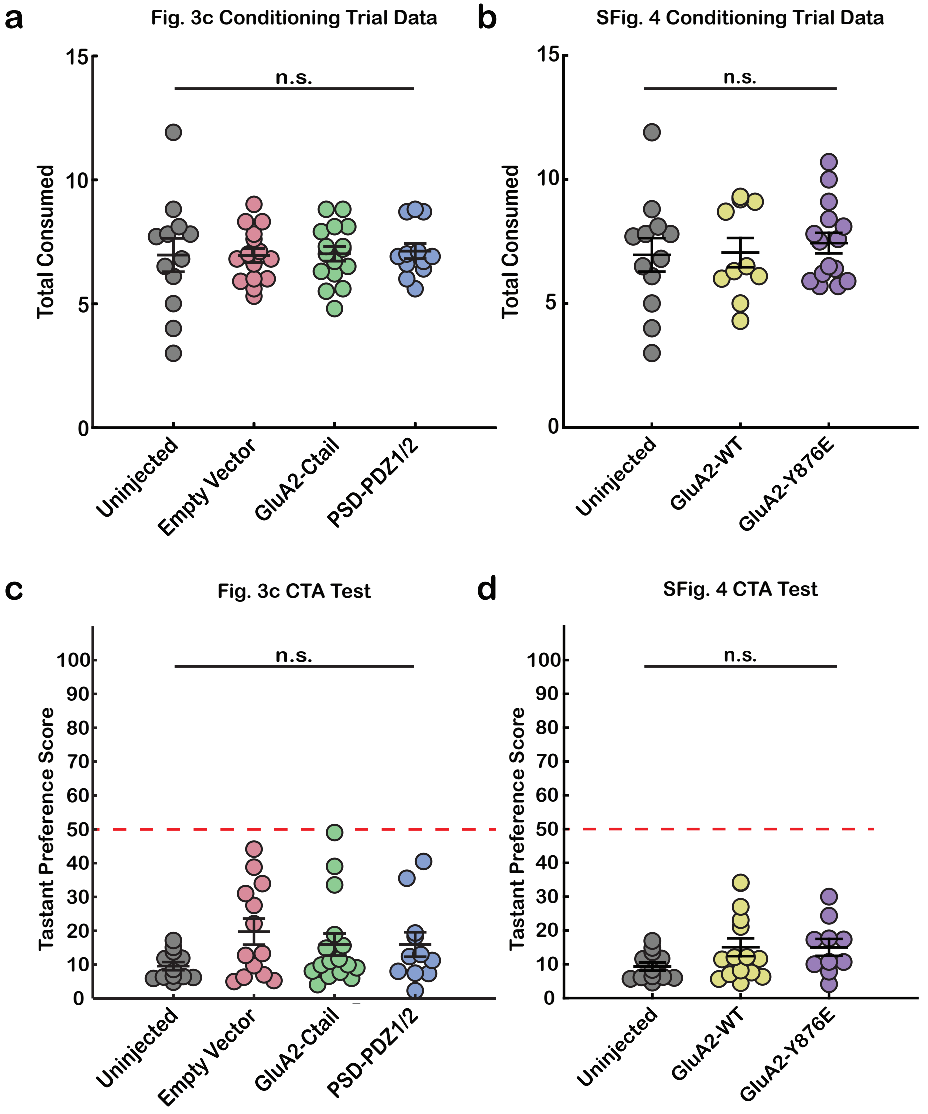

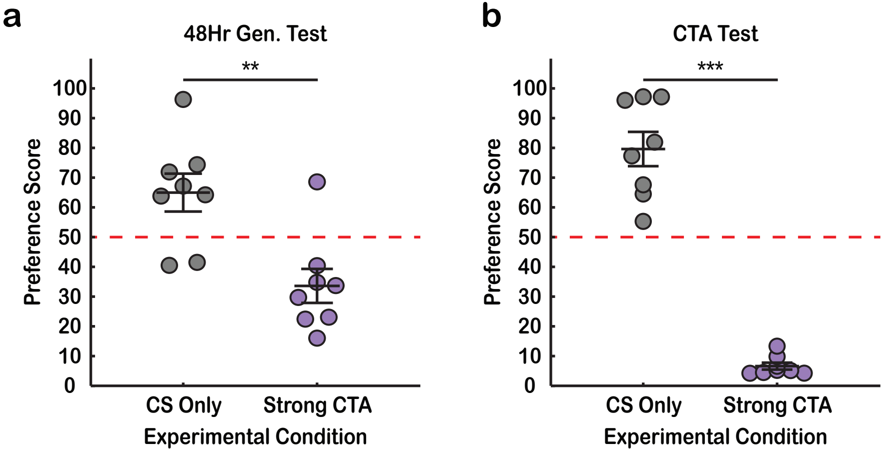

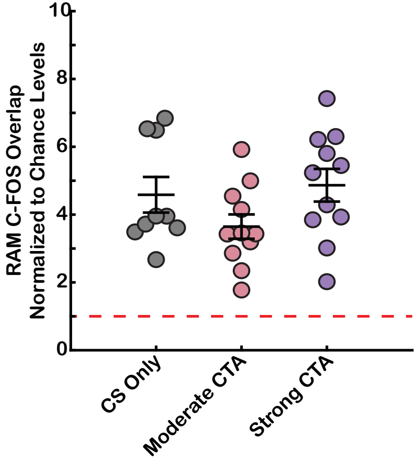
