## Supplemental Figure Legends for "Homeostatic Synaptic Scaling Establishes the Specificity of an Associative Memory"

**SFig. 1 | No sex-differences in the expression of generalized or specific CTA a)** Figure 1d data grouped by sex. No sex differences are observed in the expression of a generalized aversion. **b)** Figure 1d CTA test results grouped by sex. No sex differences are observed in the expression of CTA.

**SFig. 2 | Figure 1 supplementary behavior data a)** Figure 1b conditioning trial data (Two-sample t-test, p = 0.9215; error bars represent SEM). **b)** Figure 1c conditioning trial data (Two-sample t-test, p = 0.4040; error bars represent SEM). **c)** Figure 1d conditioning trial data (Kruskal-Wallis, p = 0.7942; Bonferroni post-hoc test, CS Only VS MCTA p = 1.000, CS Only VS SCTA p = 1.000, MCTA VS SCTA p = 1.000; error bars represent SEM). **d)** Figure 1c CTA test results (Kruskal-Wallis, p = 4.3672e-06; Bonferroni post-hoc test, CS Only VS MCTA p = 0.0000, CS Only VS US Only p = 1.000, US Only VS MCTA p = 0.0001; error bars represent SEM). **e)** Figure 1d CTA test results (One-way ANOVA, p = 4.4921e-21; Tukey-Kramer post-hoc test, CS Only VS MCTA p = 0.0000, CS Only VS SCTA p = 0.0000, CS Only VS US Only p = 0.9981, MCTA VS SCTA p = 0.6900, MCTA VS US Only p = 0.0000, SCTA VS US Only p = 0.0000; error bars represent SEM).

**SFig. 3 | Acute application of CNO activates hM4D(Gi) in brain slice a)** Resting membrane potential (V_r_) of hM4D(Gi)^+^ neuron recorded while in current clamp mode (I = 0) during a 10-minute CNO wash on. The approximate turn-over time for solution in the bath was 1-minute. Grey bars represent analysis window for V_r_. **b)** Average V_r_ during the 1^st^ and 10^th^ minute of the CNO application (n = 12; paired t-test, p = 2.8005e-04). **c)** Input resistance recorded before and after CNO treatment (n = 11; paired t-test, p = 6.8156e-05). **d)** Representative recording during 100 pA I-step before (black) and after (pink) CNO treatment. **e)** Mean firing rate VS current injection (F-I) for neurons before and after CNO treatment (n = 11; Two-Way Repeated Measures ANOVA, treatment-interaction p = 0.0299).

**SFig. 4 | Supplementary behavior data for unpaired reverse conditioning a)** Basic experimental timeline. **b)** Water trial, total consumption (Two-sample t-test, p = 0.7002). **c)** Saccharin trial, total consumption (Two-sample t-test, p = 0.0658). **d)** Tastant preference score, CTA test (Wilcoxon Ranksum, p = 0.7587).

**SFig. 5 | GluA2-Y876E extends CTA-induced generalized aversion. a)** Tastant preference scores, Gen. test 24Hrs post-conditioning (Uninjected, n = 12; GluA2-WT, n = 10; GluA2-YE, n = 15; One-Way ANOVA, p = 0.0426; Tukey-Kramer post-hoc test, UnInj. VS WT p = 0.5274, UnInj. VS YE p = 0.0340, WT VS YE p = 0.3815; error bars represent SEM). **b,c)** Gen. test 24 & 48Hrs post-conditioning (GluA2-WT paired t-test, p = 0.9431; GluA2-YE paired t-test, p = 0.0081)

**SFig. 6 | Supplementary behavior data for synaptic scaling manipulations. a)** Figure 3c conditioning trial data (One-Way ANOVA, p = 0.9904). **b)** SFigure 4 conditioning trial data (One-Way ANOVA, p = 0.7959). **c)** Figure 3c CTA test results (Kruskal-Wallis, p = 0.1950) **d)** SFigure 4 CTA test results (Kruskal-Wallis, p = 0.2050).

**SFig. 7 |** **General aversion is still present 48 hours after strong conditioning. a)** Tastant preference score from the Gen. test first conducted at 48 hours post-conditioning (CS Only, n = 8; SCTA, n = 8; Wilcoxon Ranksum, p = 0.0030). **b**) Tastant preference score for CTA following the completion of general aversion testing (Wilcoxon Ranksum, p = 1.5540e-04).

**SFig. 8 |** **The overlap of RAM-positive and c-FOS-positive cells are above chance level across experimental groups. a)** The reactivation of conditioning-active neurons (RAM^+^c-FOS^+^/DAPI^+^) was normalized to chance overlap in each group (one-sample t-test against chance level: CS Only, p < 0.0001, MCTA, p < 0.0001, SCTA, p < 0.0001).
